# Overcoming the effect of ASH1L haploinsufficiency on stem cells amenability to genome editing and differentiation into the neuronal lineage – a technical report

**DOI:** 10.1101/2021.09.27.461943

**Authors:** Seon Hye Cheon, Foster D. Ritchie, Janay M. Vacharasin, Nicholas Marinelli, Collin Cheatle, Mikayla M. McCord, Kaitlin Cox, Sofia B. Lizarraga

**Author notes:** These authors contributed equally to this work (Seon Hye Cheon, Foster D. Ritchie).

## Abstract

Genome editing and neuronal differentiation protocols have proliferated in the last decade. Mutations in genes that control pluripotency could lead to a potential obstacle with regards to the survival and differentiation potential of the genome-edited cell lines. Here we describe a protocol for the generation, and differentiation, of cell lines containing CRISPR/Cas9 induced mutations in the histone methyl transferase ASH1L. This chromatin modifier was previously implicated in hematopoietic stem cell pluripotency and is a major genetic risk factor for autism spectrum disorders (ASD). We find that haploinsufficiency of ASH1L leads to decreased NANOG gene expression leading to reduce cell survival and increased spontaneous differentiation. We report a method that provides improved single-cell survival with higher colony formation efficiency in ASH1L mutant stem cells. Additionally, we describe a modified dual-SMAD inhibition neuronal induction methodology that permits the successful generation of human neurons with mutations in ASH1L, in a smaller scale than previously reported methods. With our modified CRISPR-genome editing and neuronal differentiation protocols, it is possible to generate genome-edited stem cells containing mutations in genes that impact pluripotency and could affect subsequent cell lineage specific differentiation. Our detailed technical report presents cost-effective strategies that will benefit researchers focusing on both translational and basic science using stem cell experimental systems.

## INTRODUCTION

CRISPR-Cas9 genome editing technology is widely used for the establishment of disease models [1,2]. A D10A mutant nickase version of Cas9 (Cas9n) is only able to cut a single DNA strand [3]. Therefore, Cas9n requires a pair of offset synthetic single-guide RNAs (sgRNA) complementary to opposite strands of the target DNA region to create a double strand break [3]. The Cas9n or double nickase based CRISPR method reduces off-target effects, as having a pair of offset sgRNAs allows for greater targeting specificity [3].

One factor that needs to be considered when designing a CRISPR-knockdown model is the role of the target gene in maintaining pluripotency. Chromatin regulators play a major role in the maintenance of stem cell pluripotency [4]. Polycomb group (PcG) genes, a family of chromatin regulators, control pluripotency through suppression of developmental pathways [4]. In particular, the Polycomb Repressor Complex 2 (PRC2) is a major orchestrator of early embryonic development, and its function is carefully regulated to ensure maintenance of pluripotency [5,6]. PRC2 catalyzes the trimethylation of lysine 27 on histone H3 (H3K27me3) [7] to regulate gene expression. H3K27me3 is a repressive histone modification that is associated with silenced genes. Loss of H3K27me3 coincides with differentiation of embryonic stem cells (ESCs) [8]. Furthermore, mutations in SUZ12 and EED, two PRC2 subunits, lead to increased expression of differentiation-specific genes [8–10]. A major epigenetic mechanism that modulates development is the control of gene expression by the counteracting activities of Polycomb and the Trithorax group of proteins [11]. A core Trithorax protein, the histone methyltransferase Mixed-lineage leukemia 1 (MLL1) protein, is essential for maintenance of a primed pluripotency state in epiblast ESCs [12,13]. Loss of MLL1 leads to the unchecked redistribution of monomethylated lysine 4 on histone H3 (H3K4me1) across the genome, inducing a naïve pluripotency state in epiblast ESCs [12–14]. Together these studies suggest that editing chromatin regulatory genes associated with pluripotency could provide a unique obstacle in the development of genome edited stem cell experimental systems. Pluripotency defects could make the formation, and expansion, of single cell clones difficult. Additionally, the spontaneous differentiation of mutant ESCs makes differentiation to a specific cell lineage challenging, especially when differentiation protocols require large number of undifferentiated ESCs for the initial induction step.

Here, we describe a method for developing genome-edited ESCs that cause the downregulation of Absent, Small, or Homeotic 1-like (ASH1L) using CRISPR-Cas9 double nickase. *ASH1L* is a major genetic risk factor for ASD [15–17] and has been associated with other non-neuronal disorders like Leukemia [18]. *ASH1L* encodes a histone methyltransferase that di-methylates histone H3 on lysine 36 (H3K36me2) [19,20] and has been associated with tri-methylation of histone H3 on lysine 4 (H3K4me3) [21]. Histone mark H3K4me3 is instrumental in the maintenance of cell fate decisions of human ESCs [22]. ASH1L is proposed to counteract the repressive activity of PRC2 by preventing the tri-methylation of H3K27 [23–25]. Defects in PRC2 function led to increased differentiation [26]. ASH1L was shown to regulate cell fate specification of multipotent stem cells that include hematopoietic and mesenchymal stem cells [27,28]. Decreased quiescence and diminished self-renewal of adult hematopoietic stem cells were identified in ASH1L mutant mice [28]. In mesenchymal stem cells, ASH1L is involved in promoting osteogenesis through repression of adipogenesis [27]. Taken together, these studies suggest that perturbing the function of ASH1L in ESCs could provide a challenge when inducing mutations by genome editing.

We found that ESCs with mutations in ASH1L have variable maintenance of pluripotency, with decreased NANOG expression and increased spontaneous differentiation, causing a significant reduction in single cell clone survival during the genome editing procedure. Our methodology improves the maintenance of genome-edited ESCs leading to increased colony formation and survival of singlecell clones. In addition, we present a modified dual-SMAD inhibition protocol [29,30] for cortical neuronal differentiation that allowed us to circumvent the challenge of spontaneous differentiation of ASH1L mutant ESCs. Our method does not require the large amounts of ESCs needed for the standard dual-SMAD inhibition neural induction protocol [30,31], while still being able to successfully produce neurons. In summary, our work provides a detailed and cost-effective protocol to increase cell survival during the generation of genome edited clones with mutations in genes like ASH1L which could compromise cell survival and pluripotency maintenance

## METHODS

### Equipment, Reagents, Buffers And Culture Media Information

All the information concerning equipment, reagents, buffers, and cell culture media information is described in detail in the supplementary materials (Supplementary Tables S1 - S4).

### Designing And Cloning Of Sgrnas Targeting ASH1L

#### SgRNA Design

Four sets of sgRNAs targeting Exon 3 of ASH1L were designed using CHOPCHOP (https://chopchop.cbu.uib.no/) and Sanger (https://wge.stemcell.sanger.ac.uk/find_crisprs) (Table 1). sgRNAs were chosen using the following criterion: (1) low off-target value and (2) a difference of 5-20 base pairs (bp) between offset sgRNAs of each set. Four sets of sgRNAs were cloned into a vector expressing Cas9D10A and puromycin resistance [3]. Exon 3 is the largest exon in ASH1L with a size of 4,564bp. We designed SgRNA Sets 1 and 2 to target a region in exon 3 located ~ 3,400 bp away from the region targeted by SgRNA Sets 3 and 4 in exon 3 (Table 1 and Fig. 1A).

**Figure 1.**
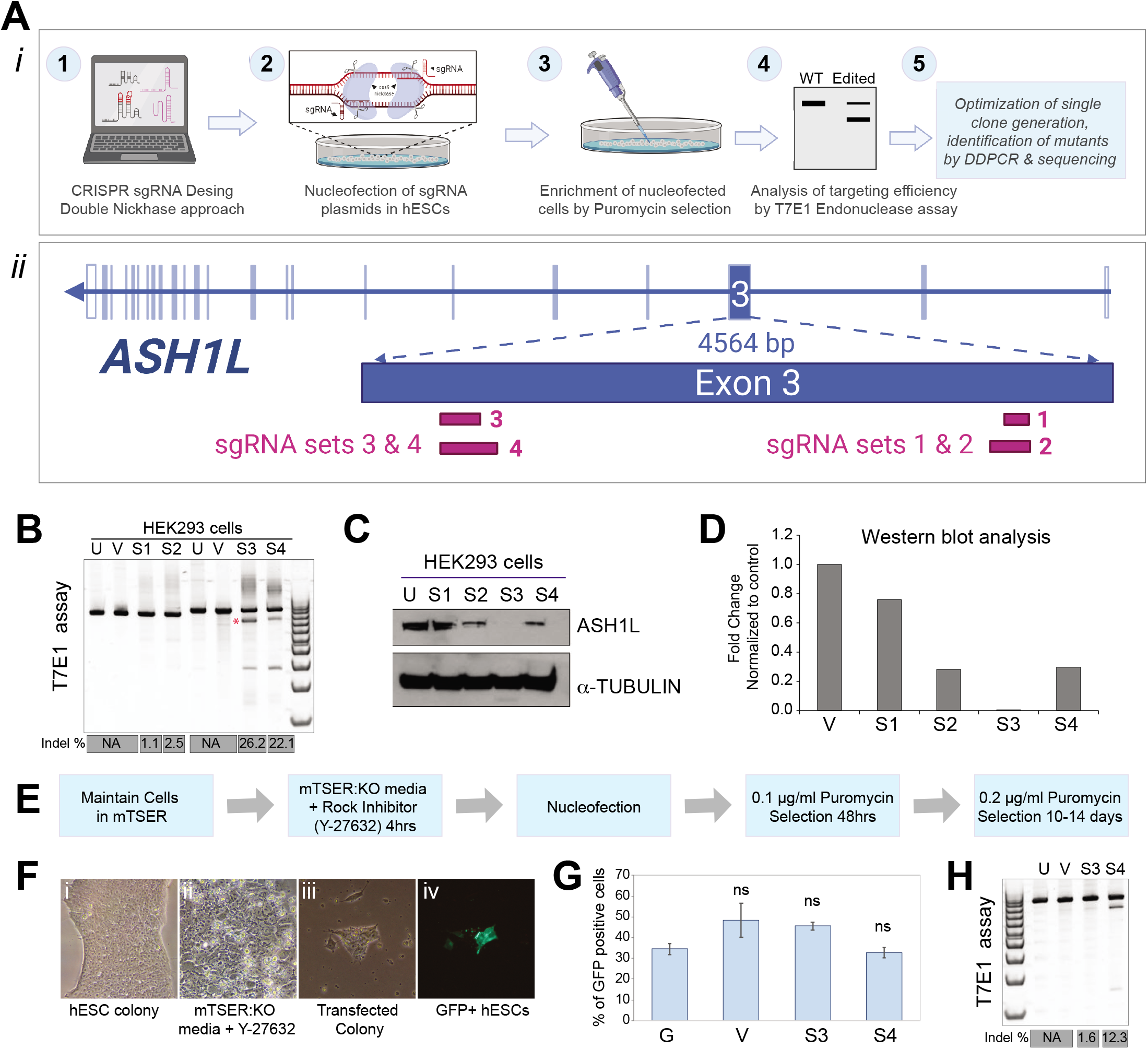
Efficient targeting of ASH1L by sgRNA constructs in HEK293 and human ESCs. (**A**) Experimental design utilizing CRISPR double Nickase approach (*i*), and diagram showing targeting strategy for exon 3 in *ASH1L* (*ii*). The areas targeted by each sgRNA set are represented by dark violet bars. SgRNAs for set 1 and 2 are located early in the exon. SgRNAs for sets 3 and 4 are located over 4000 bp downstream from the start of the exon (**B**) Analysis of sgRNA targeting efficiency by T7 Endonuclease I mismatch assay. A representative experiment is shown. Four pairs of offset complementary sgRNAs targeting ASH1L Exon 3 were designed and tested. HEK293 cells were either untreated (U) or transfected with either Px462.v2 vector (V), ASH1L-sgRNA-set1 (S1), ASH1L-sgRNA-set2 (S2), ASH1L-sgRNA-set3 (S3), or ASH1L-sgRNA-set3 (S4). SgRNA sets 1 and 2 target a different region of ASH1L Exon 3 than sgRNA sets 3 and 4, leading to different expected sizes of cleaved and uncleaved bands. SgRNA sets 3 and 4 show two prominent cleaved bands of distinct sizes. Indel percentage was calculated for all sgRNA transfected samples for the larger size cleaved band marked by a red asterisk. S1 = 1.05%; S2 = 2.49%; S3 = 26.24%; S4 = 22.1%. (**C**) Western Blot of HEK293 cells with induced mutations in ASH1L. Membranes were probed with antibodies against ASH1L and *α*-Tubulin (loading control). (**D**) Analysis of ASH1L protein levels. Data for a representative experiment is shown as fold-change normalized to control sample (V). S1 = 0.7594; S2 = 0.2807; S3 =0.0022; S4 = 0.2975. (**E**) Experimental design of ESC nucleofection and selection of transfected cells as a mix population. (**F**) Representative images of ESCs throughout the experiment. Representative images are shown for: H1 ESCs maintained with mTeSR (i); ESC colony maintained with media containing mTeSR, KO-DMEM, and ROCK inhibitor (ii); transfected colony after three days of Puromycin selection (iii); and GFP positive cells in same transfected colony (iv). (**G**) Analysis of nucleofection efficiency. Data is shown as percentage of GFP positive ESCS that were either transfected with GFP alone (G), or GFP plus either Px462.v2 vector (V), ASH1L-sgRNA-set3 (S3) or ASH1L-sgRNA-set4 (S4). GFP alone = 34.51 ± 2.61, *n* = 226 cells; GFP+ Px462.v2 vector = 48.28 ± 8.2, *n* = 174 cells; GFP + ASH1L-sgRNA-set3 = 45.54 ± 1.86, *n* = 314 cells; GFP + ASH1L-sgRNA-set4 = 32.8 ± 2.37, *n* = 314 cells. (**H**) T7 Endonuclease assay for mismatch analysis of sgRNA targeting efficiency. ESCs were either Untreated (U) or transfected with either Px462.v2 vector (V), ASH1L-sgRNA-set3 (S3), or ASH1L-sgRNA-set4 (S4). A representative experiment is shown in which the cleaved band of largest size was analyzed to calculate INDEL percentages of 1.67% (S3) and 12.26% (S4).

**TABLE-1.**
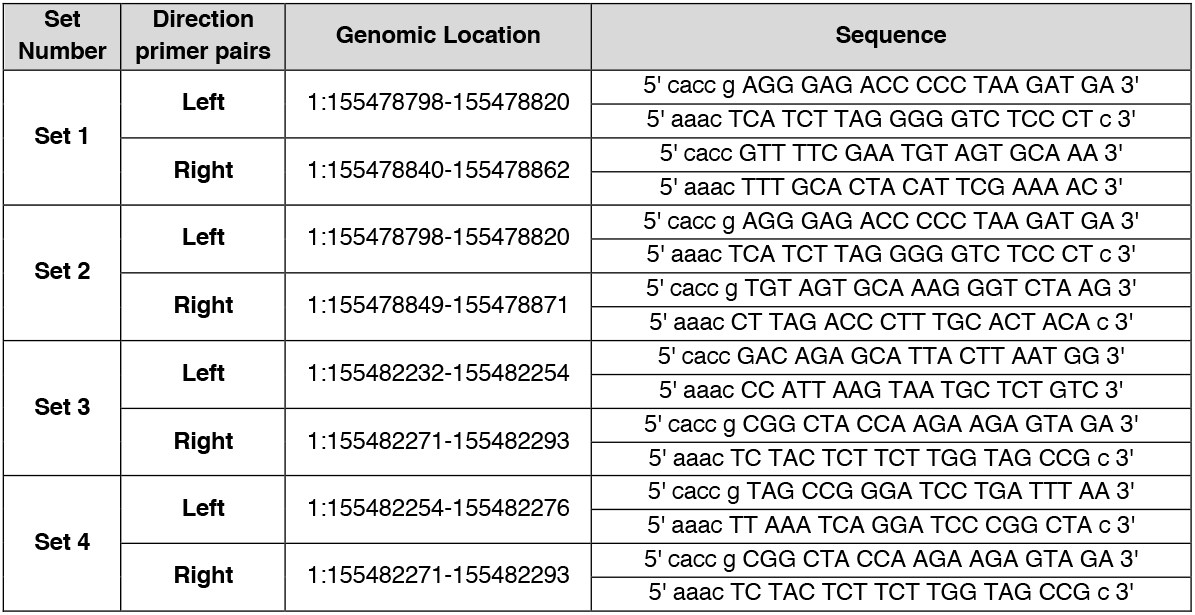
SgRNAs designed to target exon 3 in ASH1L by Double Nickase Approach. Description of sgRNA annealing primers, showing respective genomic location and sequences for each set. Genomic sequences are designated by upper case letters. Restriction cloning sites sequences are designated by lower case letters.

#### Annealing of sgRNAs duplexes

1*μ*l of each complementary sgRNA per set (100uM) were combined with 1*μ*l of 10x T4 PNK buffer, 6.5*μ*l distilled deionized water (ddH_2_0), and 0.5*μ*l T4 PNK. Oligonucleotides were annealed using a thermocycler under the following conditions: 37°C for 30 minutes, 95°C for 5 minutes, and brought down from 95°C to 25°C at 5°C/minute. Annealed oligonucleotides can be stored at −20°C for over a month.

#### Single-step digestion and ligation of sgRNAs into Px462v.2 vector

To generate CRISPR-Cas9 targeting constructs we used the pSpCas9n(BB)-2A-Puro V2.0 (Px462v.2, a gift from Feng Zhang, Addgene plasmid #62987). Px462v.2 plasmid [3] was simultaneously digested and ligated to annealed oligo duplexes diluted 1:200. Reactions in a final volume of 20μl contained 2*μ*l of the diluted duplex 100ng of Px462v.2, 2*μ*l of Tango buffer 10x, 1*μ*l DTT (1mM), 1*μ*l ATP (1mM), 1*μ*l FastDigest BbsI (Bpi), 1*μ*l T4 DNA ligase, and ddH_2_O up. The reaction was conducted in a thermocycler (37°C for 5 minutes, 23°C for 5 minutes for 6 cycles). Plasmid-Safe Exonuclease (PSE) was used to remove unwanted recombination products, using 1*μ*l of the ligation reaction product, 1.5*μ*l of 10X plasmid safe Buffer, 1.5*μ*l of ATP (10mM), and 1*μ*l of PSE. This reaction was incubated at 37°C for 30 minutes.

### Human Embryonic Kidney (HEK) 293 and Stem Cell Tissue Culture Conditions

#### HEK293 cultures

HEK293 cells were grown in DMEM media containing FBS, L-glutamine and Pen/Strep and were passaged every 3 to 4 days (Supplementary Table S1).

#### Stem Cell Culture

H1ESCs (WI-cell catalog #WA01) [32] were grown in Primaria 6-well plates coated with Matrigel (extracellular matrix substrate). ESCs were fed daily with mTeSR (2ml), and were inspected daily for spontaneous differentiated colonies that were manually removed. Removal of differentiating colonies ensures the pluripotency of the cultures. Cells were passaged at a 1:6 ratio every 4 to 7 days with ReLeSR. Cultures were tested monthly for mycoplasma.

### sgRNA Introduction Into Mammalian Cells

#### Transfection of HEK cells

HEK cells were seeded at 1×10e5 cells per well. 70% confluent cultures were transfected using Lipofectamine 2000 Transfection Kit per manufacturer’s protocol.

#### Nucleofection of Human ESCs

Prior to dissociation, cells were incubated with a 1:1 mixture media containing mTeSR + KO-DMEM + 10*μ*M ROCK inhibitor (mKOR) for 4 hours. Cells were incubated in 1 ml of gentle cell dissociation reagent (GCDR) for 10 min at 37°C. Cells in GCDR were dis-lodged using 2ml of mKOR and the remaining cells were harvested in an additional 4ml of mKOR per well. Harvested cells were centrifuged at 100g for 5 minutes. ESCs were gently single cell dissociated with a 1 ml pipette tip. For a final 20*μ*l reaction, 3-5×10^5^ cells were combined with 16.4*μ*l P3 solution, 3.6*μ*l P3 supplement and DNA solution (pmax-GFP 100ng, Px462v2-sgRNA-Left 200ng, Px462v2-sgRNA-Right 200ng). Cell mixture was gently loaded onto a cuvette strip to avoid air bubbles. Cells were nucleofected using CA-137-high efficiency program on 4D Amaxa nucleofection unit. After nucleofection, cells were incubated at room temperature (RT) for 10 minutes. Cells were then gently resuspended in 80*μ*l of pre-warmed mTeSR + 10*μ*M ROCK inhibitor and plated on a Matrigel-coated Primaria 24-well plate. Transfected cells were selected with 0.2*μ*g/ml puromycin for 10-14 days after nucleofection.

### Identification Of Targeting Efficiency By Mismatch Analysis Using T7 Endonuclease 1 (T7E1)

#### Genomic DNA Extraction from HEK cells and ES Cells

We extracted genomic DNA from transfected HEK cells using the QuickExtract™ DNA extraction solution. Genomic DNA was diluted to 100ng/*μ*l for genomic PCR reactions.

#### PCR amplification of target region

To ensure fidelity during PCR amplification we used Herculase-II Polymerase (1*μ*l) in combination with genomic DNA (1*μ*l), 5X Herculase buffer (10*μ*l), 100mM dNTP (1*μ*l), 0-2*μ*l of MgCl2 (depending on GC content), 1*μ*l each of forward and reverse primers (10*μ*M), and ddH_2_O to a final volume of 50*μ*l. Exon 3 primers (Supplementary Table S5) were designed to generate ~ 900 bp amplicons containing regions targeted by the different sgRNA sets. Reaction conditions were 95°C for 5 min, 30-35 cycles of [95°C for 30 seconds, 58-62°C for 30 seconds, 72°C for 30 seconds per 1kb], followed by 72°C for 5 minutes, and hold at 4°C. The PCR product was run on a 2% TAE agarose gel for 1 −1.5 hours at 150V. Gel bands were extracted using the Qiagen QIAquick Gel Extraction Kit.

#### T7E1 Assay

Reactions containing 19*μ*l of a heteroduplex and 1*μ*l of T7E1 were run on a 4 −20% Novex TBE gel at 200V for 30 minutes. Gels were stained with SyBr Gold at RT for 30 minutes and were imaged using a Chemidoc MP Imaging System (Bio-Rad). Percent modification and the fraction cleaved by T7E1 were calculated as previously described [33] using the following formulas:

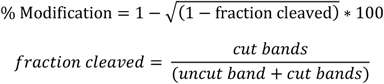

### Generation Of Single-Cell Clones From ESCs

Single cell clones were generated 10 to 14 days after nucleofection by dissociating ESCs colonies that were selected with 0.2*μ*g/ml of puromycin. ESCs were dissociated with GCDR (1 ml) for 10 minutes at 37°C, harvested in mKOR (4ml) and centrifuged at 100g (5 minutes). Cell pellets were resuspended, single cell dissociated, and plated on a 96-well Matrigel coated plate at 1 cell per well in mTeSR +10*μ*M ROCK inhibitor + 10% CloneR. After 24 hours the media was changed to mTeSR + 10*μ*M ROCK inhibitor + 2.5% CloneR.

### Digital Droplet PCR (ddPCR) Screening For Mutations

To efficiently screen for mutant clones, we used a digital droplet PCR (ddPCR) approach using genomic DNA [34–36]. Sequence specific primer pairs were designed for amplification of the CRISPR/Cas9 targeted region (Supplementary Table S5). Additionally, FAM and HEX probes were designed to target wildtype and mutant regions, respectively [34,37]. Genomic DNA was extracted from the mutant ESCs using the DNeasy Blood and Tissue Kit. To ease amplification of target regions, we digested the genomic DNA (50ng) with EcoRI-HF at 37°C for 15 minutes, following manufacturers protocol. Oil droplets generated using a DG8 cartridge were transferred to a Bio-Rad 96-well plate that was then sealed using a pierceable foil seal. Target amplification was done using the following conditions: 95°C for 10 minutes, 40 cycles of [94°C for 30 seconds, 60°C for 1 minute], 98°C for 10 minutes, and 4°C hold. The reaction was analyzed using a QX200 Droplet Reader, equipped with QuantaSoft software as described [35–37].

### Protein Extraction, Quantification and Western Blot

#### Protein Extraction and Quantification

Cells were placed on ice, washed with ice cold sterile PBS (1X), and lysed in RIPA buffer containing protease and phosphatase inhibitors (1:500 each) (70*μ*l per well for 12-well plates). Cells in RIPA buffer were collected, gently vortexed, and incubated on ice for 10 minutes. To remove cell debris, lysates were centrifuged at 14,000rpm for 10 minutes at 4°C. Collected supernatant was flash frozen in liquid nitrogen and stored at −80°C till further use. Protein concentrations were estimated using the Pierce BCA protein_assay kit per manufacturers protocol using a Spectramax i3 plate reader.

#### Western Blot analysis

10*μ*g of total protein were loaded per sample in a pre-cast Bolt 4-12% Bis-Tris Plus gel and run at 200V for 25 minutes. Samples were transferred on ice using Towbin buffer for transfer of high molecular weight proteins at 41V, 0.45A, and 19W for 90 minutes. Blots were processed as previously described [38] using primary antibodies for ASH1L and GAPDH (Supplementary Table S3) diluted in 3% BSA/TBST, and HRP conjugated secondaries diluted (1:15,000) in 5% milk/TBST. Blots were probed with ECL for 5 minutes, and developed using CL-XPosure™ Film on a Konica SRX-101A Film Processor.

### Sequencing And Analysis

A 972 base pair amplicon covering the CRISPR-Cas9 targeted region of ASH1L in exon 3 was amplified using site specific primers, and was sequenced with forward and reverse primers using standard Sanger sequencing (Supplementary Table S5). DNA sequences were analyzed using Tracking of Indels by Decomposition (TIDE) [39] and CRISPR-ID [40].

### Quantification Of Gene Expression By Quantitative PCR (qPCR)

RNA was extracted using the RNeasy Micro kit following the manufacturer’s instructions. RNA was converted to cDNA using the Sensifast cDNA conversion kit following the manufacturer’s instructions. qPCR was performed using the intercalating dye SYBR green on the CFX96 Touch Real Time PCR Detection System (Bio-Rad). qPCR primers were designed to identify all isoform variants and were validated by IDT (Supplementary Table S3). Three technical replicates were used per sample using 4ng of total cDNA and 1*μ*M of qPCR primers per reaction, under the following conditions: 95°C for 10 minutes, followed by 40 cycles of 95°C for 15 seconds and 60°C for 1 minute. Ct values were generated by the Bio-Rad software, and relative gene expression with respect to control was generated by the CFX Manager Software.

### Small-Scale Neuronal Induction Of ASH1L Genome Edited Stem Cell Lines

Cells were seeded in one well of a 24-well Primaria plate with 5 to 8 x 10^5^ cells per well and placed in *Stemdiff* neuronal induction medium containing ROCK Inhibitor for 24 hours. Cells were then switched to Neural Induction Media (NIM) (Supplementary Table S1) to elicit a dual-SMAD inhibition [29]. Cells were maintained with daily media changes for 11 days. At day 12, cells were expanded using GCDR at 1:1 to a 24-well plate and continued to be maintained in NIM. From day 17 to day 20, cells were maintained in NMM containing FGF (20 ng/ml), to induce proliferation of neuronal progenitors. On day 21 wells containing cells with increased differentiation of non-neuronal cells were cleaned by placing harvested cells were placed for 5 minutes at 37°C in plate pre-coated with poly-ornithine (2hrs to overnight at 37°C). Cleaned cells were expanded 1:2 and maintained with NMM every other day with half media changes or frozen at 2×10e6 cells/ml and stored in liquid nitrogen [29].

## RESULTS

### Targeting Of ASH1L In HEK293 and Human ESCs

Since HEK293 cells are easier to transfect, we first tested targeting efficiency of chosen sgRNAs to ASH1L gene in this cell type. Mutant ASH1L HEK293 cell lines were generated using the CRISPR double nickase approach and targeting 2 different regions in exon 3 (Fig. 1A). This approach uses Cas9n which can only cut one DNA strand [3], requiring two offset sgRNAs (sgRNA set) that will direct Cas9n to target two different sites within 10 to 30 bp apart in each DNA strand (Ran *et al*., 2013) (Table 1). Therefore, a significant reduction in off-target effects is observed with the double nickase approach.

Mutant ASH1L HEK293 lines were confirmed using a T7 Endonuclease 1 (T7E1) Assay that cleaves mismatched DNA (Fig. 1B). sgRNA Sets 1 and 2 target regions in exon 3 that are 3,400 bp apart from the target regions of sgRNA Sets 3 and 4. Therefore, it is possible that cleaved and uncleaved bands will have different sizes depending on what region is analyzed. After quantification of the percent modification, we found that ASH1L was most efficiently targeted by sgRNA Sets 3 and 4 (Table 1 and Fig. 1B). Induced mutations in ASH1L impacted the protein levels of ASH1L, most significantly decreased in cells nucleofected with sgRNA Sets 2, 3 and 4 (Fig. 1C-D). A reduction in ASH1L protein levels was expected from our analysis of targeting efficiency for sgRNA sets 3 and 4. However, the reduced ASH1L levels associated with sgRNA set 2 were unexpected and could be indicative of mutations introduced due to random repair in ASH1L in regions that fall outside of the amplicon area used for the T7E1 assay. Given that sgRNA Sets 3 and 4 targeted ASH1L most efficiently showing the greatest reduction in ASH1L protein levels, we used Sets 3 and 4 for subsequent experiments in ESCs.

ESCs grow as tight and compact colonies that are difficult to transfect at high efficiency with regular transfection protocols (Fig. 1E-F), requiring single cell dissociation for more efficient transfection. However, single cell dissociation of ESCs presents an additional challenge as single cell ESCs are prone to increased cell death and differentiation. To overcome these challenges, we incubated cells with mKOR (mTeSR + KO-DMEM + 10*μ*M ROCK inhibitor) for 4 hours prior to dissociation. ROCK inhibitor increases ESC survival by preventing dissociation-induced apoptosis [41]. Cells incubated with mKOR media showed a loosened colony structure (Fig. 1E-F). ESCs dissociated into single cells were nucleofected with a GFP plasmid plus either Px462.V2 empty vector or Px462.V2 containing the sgRNA plasmid sets. Analysis of the percentage of GFP+ cells showed no significant difference in the efficiency of transfection between control or ASH1L sgRNA groups (Fig. 1G). Mutant ASH1L ESCs were then confirmed using the T7E1 Assay (Fig. 1H). Quantification of the T7E1 Assay showed greater targeting efficiency in cells nucleofected with the sgRNA-set4 plasmids (Fig. 1H). It is important to note that sgRNA targeting efficiency is not necessarily consistent across different cell types. This could be due to variations in Single Nucleotide Polymorphisms or differences in the chromatin states in the targeting regions of different cell types [42,43]. Therefore, for genome editing experiments in ESCs we used the Set 4 sgRNAs given its improved targeting efficiency in ESCs.

### Optimization Of Single Cell Survival And Colony Formation In ASH1L Mutant ESCs

Nucleofected ESCs were seeded as single cell clones in a 96-well plate. However, there was no survival of cells seeded with mKOR media. To improve the survivability of single cell clones, we utilized CloneR, a supplement that increases single cell survival at low clonal densities. We cultured 32 vector alone single cell clones and 144 sgRNA-set 4 single cell clones in mTeSR + 10*μ*M ROCK inhibitor + 10% CloneR for 24 hours (Fig. 2A). After 24 hours, the media was changed to mTeSR + 10*μ*M ROCK inhibitor + 2.5% CloneR (Fig. 2A). Colony formation and survival percentage of ESCs transfected with the pX462.V2 control plasmid were 68.75% each when maintained with both CloneR and ROCK inhibitor (Fig. 2A). However, Set 4 sgRNA-nucleofected colonies showed reduced colony formation (33.33%) and survival (27.78%) (Fig. 2A). A total of 40 ASH1L sgRNA set 4 surviving clones were sub-cloned for further analysis.

**Figure 2.**
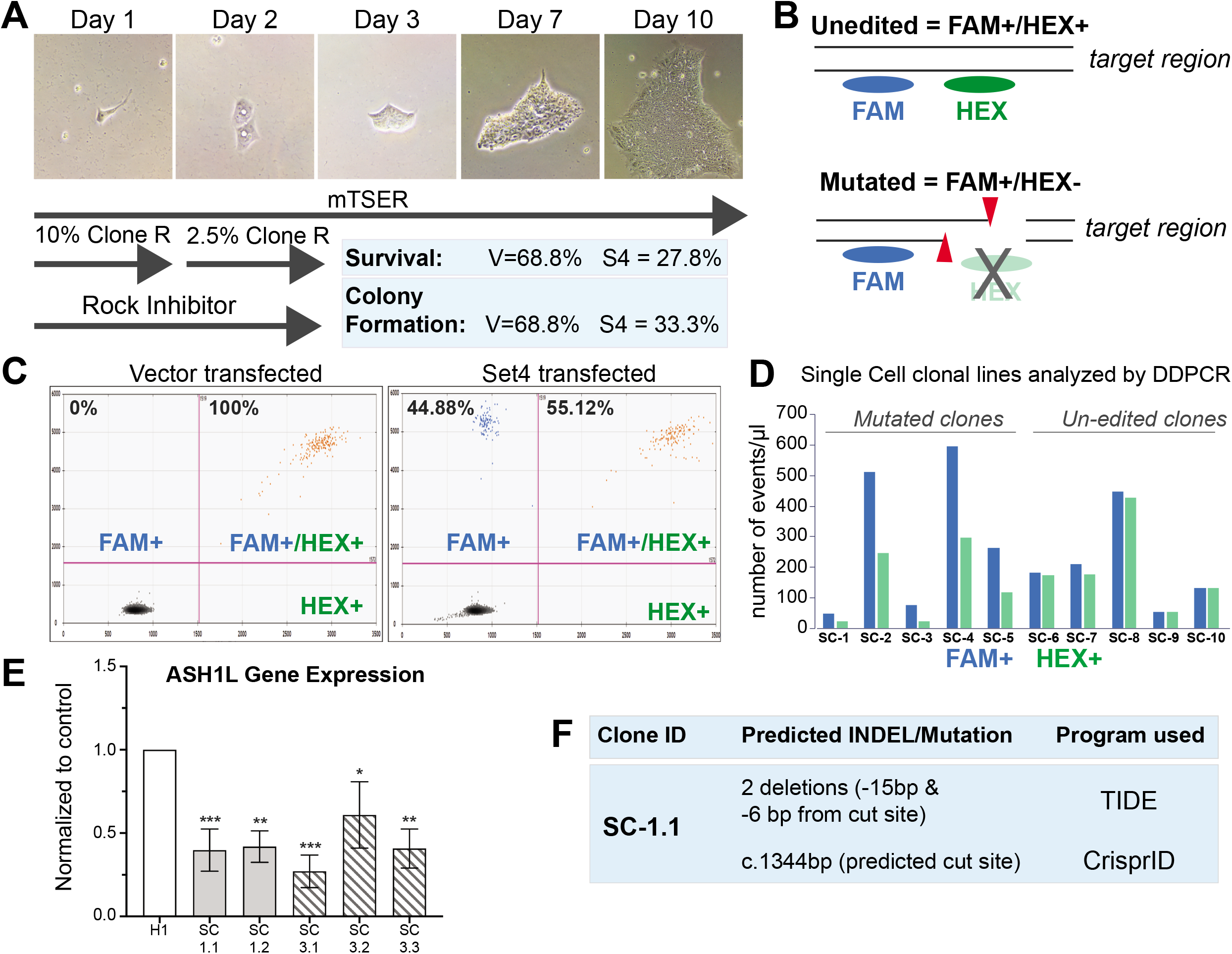
Identification and molecular characterization of ASH1L induced mutations. (**A**) ESC single-cell clonal expansion into a cell colony in a 10-day period. Transfected ESCs were dissociated as single cells and grown for 24hrs using 10% CloneR + ROCK inhibitor for 24 hours followed with 2.5% CloneR for additional 24 hours. Representative images show the longitudinal progression from a single cell to a hESC colony. Mutant colony formation and survival rates compared to control ESCs. H1 ESCs were either transfected with px462.V2 plasmid or with ASH1L-sgRNA-set4 plasmids and seeded as single cells for px462 transfected cells (32 single cells) and ASH1L-sgRNA-set4 transfected cells (144 single cells). (**B**) Diagram of ddPCR strategy for mutant identification. In the absence of any mutations, FAM and HEX probes will bind to both target regions. If a mutation is present, the HEX probe will be unable to bind and only the FAM probe will be registered. (**C**) Comparison of control and ASH1L mutant cells obtained without single cell clonal selection and analyzed by DDPCR for proof of feasibility. A 2D plot of droplet fluorescence for FAM and HEX levels is shown. Percentage of FAM+/HEX+ signal and FAM+/HEX-signal is shown for control H1 ESCs (left panel) and ASH1L ESCs transfected with sgRNA set-4 (right panel). Control H1 ESCs showed 100% FAM+/HEX+ levels. ESCs transfected with sgRNA set-4 showed 44.88% FAM+/HEX-levels and 55.12% FAM+/HEX+ levels. (**D**) Analysis of single cell clonal populations for FAM+/HEX+ and FAM+/HEX-signal. A total of 40 single cell clones obtained after nucleofection with ASH1L-sgRNA set-4 were analyzed. A subset of the single cell clones is shown as potential ASH1L mutant clones identified by ddPCR (SC-1 to SC-5) and as clones not identified as mutated or un-edited (SC-6 to SC-10). (**E**) Analysis of ASH1L gene expression: SC-1.1 = 0.3976 ± 0.1208; SC-1.2 = 0.4190 ± 0.0900; SC-3.1 = 0.2709 ± 0.0922; SC-3.2 = 0.6089 ± 0.1899; SC-3.3 = 0.4076 ± 0.1116. (**F**) Table showing different approaches to identify genome-edited mutations.

### Screening For ASH1L Mutations By ddPCR

Digital Droplet PCR (ddPCR) has emerged as a fast-screening tool to identify genome edited clones by analyzing the genomic DNA [35,36]. We designed FAM and HEX labelled probes to target the un-edited and mutated regions, respectively (Fig. 2B, Supplementary Table S5) [34,37]. We first validated the probes using a mixed population approach. ESCs were nucleofected with either Px462-V2 empty vector plasmid [3] or the ASH1L-sgRNA Set 4 plasmids. Each entire nucleofected population was plated in a single well of a 24-well plate. Genomic DNA extracted from these cells was analyzed for FAM and HEX double-positive or FAM-only positive signals. A FAM and HEX double-positive signal indicates an un-edited allele because the reference and mutant probes binds to it [35,36]. In contrast, the mutant group would only have a FAM positive signal since the HEX probe will not be able to bind if a mutation is present (Fig. 2B). Control ESCs contained 100% FAM+/HEX+ signal, while ASH1L mutant cells had 55.12% FAM+/HEX+ signal and 44.88% FAM+ only signal (Fig. 2C).

After optimization of single-cell survival, we used ddPCR to confirm the targeted mutation of the 40 surviving clones. Mutations were identified using the FAM and HEX probes to analyze genomic DNA (Fig. 2B). From the ddPCR analysis, we were able to identify 5 possible mutant clones (SC-1, SC-2, SC-3, SC-4, and SC-5) (Fig. 2D). These clones showed approximately half HEX+ droplets for each sample, compared to approximately equal FAM+ and HEX+ droplets in un-edited cell lines. These results indicate that in our mutant clones, half of the droplets screened were unable to bind the mutant probe, suggesting a mono-allele mutation [34]. Events from all 40 clones and representative 2D plots of droplet fluorescence for mutant clones are in the Supplementary Material (Supplementary Fig. S1A). To validate our approach, we analyzed ASH1L gene expression in five mutant ASH1L cell lines and found significant downregulation of ASH1L mRNA expression in all mutant ESCs (Fig. 2E). Mutations were also confirmed through sequencing of the mutant single-cell clones. Analysis of Sanger Sequencing data using TIDE and CRISPR-ID programs predicted a deletion event in ASH1L Exon 3 of SC-1.1 mutant clone (Fig. 2F) [39,40,44]. Predicted indels and corresponding aberrant sequence percentage can be found in Supplementary Material (Supplementary Fig. S2).

### Testing The Effect Of ASH1L Mutations In Pluripotency And Cell Lineage Marker Expression

ASH1L is proposed to have a role in stem cell fate and maintenance of pluripotency in hematopoietic stem cells [27,28]. We observed Increased spontaneous differentiation in ASH1L mutant ESC lines, so we quantified the mRNA expression of NANOG, a gene that is highly expressed in undifferentiated ESCs [45]. All the ASH1L subclonal mutant lines analyzed had a significant downregulation of NANOG compared to the control, suggesting potential decreased pluripotency in mutant ESC lines (Fig. 3A). Additionally, to assess the extent to which mutations in ASH1L could lead to changes in cell lineage, we analyzed the expression levels of three different cell lineage marker genes, AFP (endoderm), MyoD (mesoderm), and GFAP (neuroectoderm) (Fig. 3B). Expression levels in the mutant ESCs varied between cell lines, with a trend of higher AFP expression in SC-3.1 and SC-3.2 clones. Despite the variability observed, no statistically significant difference was observed between control and mutant ESCs for any of the cell lineage markers utilized.

**Figure 3.**
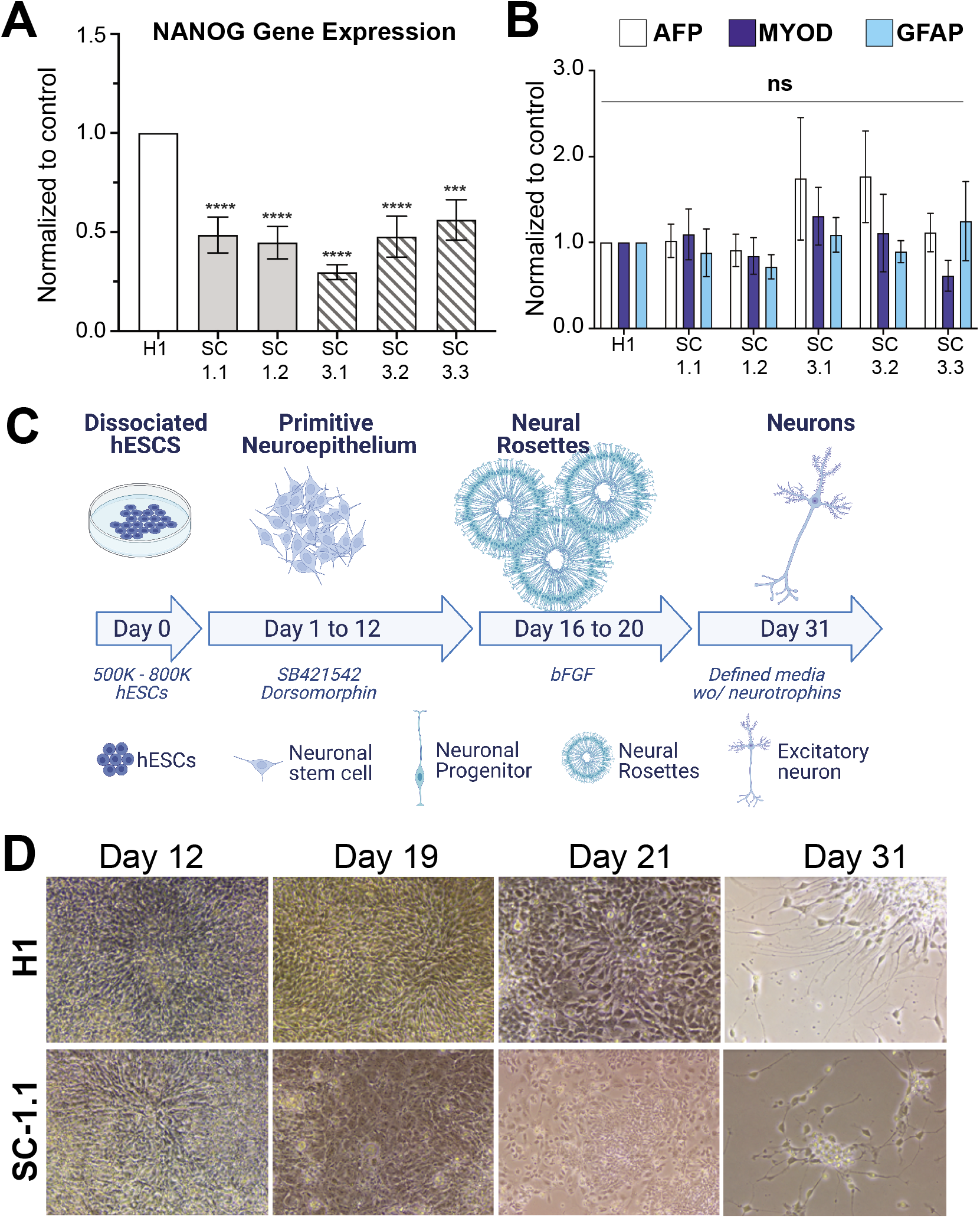
Characterization of gene expression in ASH1L genome edited lines and development of a small-scale neuronal induction protocol. (**A-B**) Analysis of gene expression for control (H1) and ASH1L mutant (SC-1.1, SC-1.2, SC-3.1, SC-3.2, SC-3.3) cell lines for pluripotency (**A**) and germ cell layer markers (**B**). (**A**) Nanog gene expression: SC-1.1 = 0.4851 ± 0.0910; SC-1.2 = 0.4463 ± 0.0817; SC-3.1 = 0.2973 ± 0.0377; SC-3.2 = 0.4764 ± 0.1034; SC-3.3 = 0.5614 ± 0.1020. (**B**) Gene expression of endoderm layer marker AFP (clear columns). SC-1.1 = 1.0185 ± 0.1934; SC-1.2 = 0.9077 ± 0.1880; SC-3.1 = 1.7402 ± 0.7136; SC-3.2 = 1.7638 ± 0.5344; SC-3.3 = 1.1139 ± 0.2223. Gene expression of mesodermal layer marker MyoD (dark blue columns). SC-1.1 = 1.0916 ± 0.2950; SC-1.2 = 0.8411 ± 0.2119; SC-3.1 = 1.3029 ± 0.3343; SC-3.2 = 1.1071 ± 0.4498; SC-3.3 = 0.6125 ± 0.1791. Expression for neuroectodermal cell lineage marker GFAP (light blue columns). SC-1.1 = 0.8792 ± 0.2759; SC-1.2 = 0.7172 ± 0.1404; SC-3.1 = 1.0862 ± 0.2012; SC-3.2 = 0.8911 ± 0.1278; SC-3.3 = 1.2447 ± 0.4589. (**A-B**) Statistical analysis was done using paired t-test, N=5 independent experiments, data is shown as mean ± standard error of the mean, **P < 0.01, *P < 0.05. (**C**) Schematic showing the small scale modified dual SMAD inhibition neuronal induction protocol. Cell types expected at different stages of neuronal induction. (**D**) Representative images of H1 WT ESCs and ASH1L genome edited line SC-1.1 at specific timepoints of the small-scale neuronal induction approach.

### Mutations In ASH1L Do Not Prevent Neuronal Differentiation

Since mutations in ASH1L have previously been associated with several neurological disorders, including ASD [15,17], Attention-deficit/hyperactivity disorder [46], and Tourette syndrome [47,48], we tested the viability of our mutant ASH1L ESCs for neuronal induction. Our lab routinely uses a dual-SMAD inhibition protocol that leads to the generation of excitatory cortical neurons [29]. This protocol generally requires between 1.5 to 2 million cells per induction [29,31]. The increased spontaneous differentiation of ASH1L mutant ESCs made it difficult to obtain the number of cells required for our standard neuronal induction protocols [29,31]. However, using a modified small-scale dual-SMAD inhibition neuronal induction, requiring between 500,000 to 800,000 cells per induction, we were able to successfully induce two ASH1L mutant ESC subclonal lines (SC-1.1 and SC-3.1) to differentiate into neurons (Fig. 3C-D, Supplementary Fig. S3). These results suggest that mutations in ASH1L are not preclusive to driving the differentiation towards the neuronal lineage.

## DISCUSSION

ASH1L regulates quiescence and self-renewal in hematopoietic stem cells [28]. Because of this, creating genome-edited ESC lines with induced mutations in ASH1L could become problematic as pluripotency of ESCs could be potentially altered. Here, we described a detailed protocol to develop genome-edited stem cell lines with mutations in ASH1L, using CRISPR-Cas9 double nickase. We strived to make this technical report as comprehensive as possible for ease of implementation.

During the establishment of single-cell ASH1L mutant clones, we observed decreased survival and increased spontaneous differentiation. The decreased survival and increased spontaneous differentiation prevented efficient colony formation from single cell clones. We first used ROCK inhibitor as it has been shown to decrease apoptosis in dissociated ESCs [41]. However, we observed no colony formation with just ROCK inhibitor. We found that media containing ROCK inhibitor with the addition of CloneR, a supplement that improves single cell colony survival, led to greater colony formation and survival, and less spontaneous differentiation of our mutant cell lines. We found a significant decrease in the expression of NANOG in all mutant ESCs. NANOG is a marker of undifferentiated ESCs. NANOG deficiency in ESCs leads to greater spontaneous differentiation towards the endoderm lineage [45]. While we found a trend in two lines showing higher expression for AFP a marker of endoderm lineage, in general we observed variable expression of cell lineage markers, but none were statistically significant.

We used ddPCR to efficiently identify mutations generated by non-homologous end joining repair mechanisms using multiple probes [34,37]. Using FAM and HEX probes designed specifically to differentiate between the non-targeted and the targeted sequences, we were able to distinguish between control and mutant alleles even in cultures that were not from a single clone. We screened for mutations across 40 different single-cell ASH1L mutant clones and found that the clones were primarily heterozygous, due to the continued presence of the HEX probe. Based on reports showing increased lethality associated with homozygous ASH1L mutations in mouse [49,50] we infer that in our system homozygous mutations might not be present due to the importance of ASH1L in cell survival. In summary, as previously reported ddPCR provides an affordable and time efficient option of detecting mutations in genome-edited cell lines

Additionally, we assessed the reliability of these cell lines for neuronal studies by differentiating them via a small-scale neuronal induction protocol. Neuronal inductions using the dual-SMAD inhibition approach typically call for 1.5 to 2 million cells per induction [29,31]. With cell lines that show high spontaneous differentiation, these number of cells are difficult to obtain. Through our small-scale neuronal induction, which requires between 500,000 to 800,000 cells per well, we can generate neurons. This approach provides a cost-effective strategy to still establish experiments using fewer cells, which could be applicable to other cell lines irrespective of the genetic background.

In summary, our work provides a guide for those having difficulties with survival of stem cells after CRIPSR-mediated genome editing. In addition, we provide a small-scale neuronal induction protocol that will allow for generation of neurons in cultures in which stem cell number are a limitation. This modified neuronal induction protocol also provides an alternative method to reduce the overall laboratory cost of the generation of human neurons in vitro, making it more accessible to all laboratories.

## ACKNOWLEDGEMENTS

The authors would like to thank the Twiss laboratory for the use of their QPCR and DDPCR equipment. We are especially grateful to Dr. Seung Joon Lee for training on the DDPCR equipment and to the laboratory of Dr. Jill Turner for training on qPCR methods. We thank Dr. Evelyn Chukwurah and Trevor Moreland for critically reading this manuscript.

## AUTHORSHIP CONFIRMATION STATEMENT

**Seon Hye Cheon:** Conceptualization, Methodology, Investigation, Data curation. **Foster D. Ritchie:** Conceptualization, Writing first draft, review & editing, Data Curation, Visualization. **Nicholas Marinelli:** Methodology and Investigation. **Janay M. Vacharasin:** Data Curation and editing the manuscript. **Mikayla McCord:** Data Curation. **Collin Cheatle:** Visualization. **Kaitlin Cox:** Visualization. **Sofia B. Lizarraga:** Conceptualization of study, Writing, review and editing, Visualization, Data Curation, Supervision, Project administration, Funding acquisition.

## REFERENCES

1. Freiermuth JL, IJ Powell-Castilla and GI Gallicano. (2018). Toward a CRISPR Picture: Use of CRISPR/Cas9 to Model Diseases in Human Stem Cells In Vitro. J Cell Biochem 119:62–68.

2. Tschaharganeh DF, SW Lowe, RJ Garippa and G Livshits. (2016). Using CRISPR/Cas to study gene function and model disease in vivo. FEBS J 283:3194–203.

3. Ran FA, PD Hsu, CY Lin, JS Gootenberg, S Konermann, AE Trevino, DA Scott, A Inoue, S Matoba, Y Zhang and F Zhang. (2013). Double nicking by RNA-guided CRISPR Cas9 for enhanced genome editing specificity. Cell 154:1380–9.

4. Lessard JA and GR Crabtree. (2010). Chromatin regulatory mechanisms in pluripotency. Annu Rev Cell Dev Biol 26:503–32.

5. Faust C, A Schumacher, B Holdener and T Magnuson. (1995). The eed mutation disrupts anterior mesoderm production in mice. Development 121:273–85.

6. Pasini D, AP Bracken, MR Jensen, E Lazzerini Denchi and K Helin. (2004). Suz12 is essential for mouse development and for EZH2 histone methyltransferase activity. EMBO J 23:4061–71.

7. Xu C, C Bian, W Yang, M Galka, H Ouyang, C Chen, W Qiu, H Liu, AE Jones, F MacKenzie, P Pan, SS Li, H Wang and J Min. (2010). Binding of different histone marks differentially regulates the activity and specificity of polycomb repressive complex 2 (PRC2). Proc Natl Acad Sci U S A 107:19266–71.

8. Chamberlain SJ, D Yee and T Magnuson. (2008). Polycomb repressive complex 2 is dispensable for maintenance of embryonic stem cell pluripotency. Stem Cells 26:1496–505.

9. Boyer LA, K Plath, J Zeitlinger, T Brambrink, LA Medeiros, TI Lee, SS Levine, M Wernig, A Tajonar, MK Ray, GW Bell, AP Otte, M Vidal, DK Gifford, RA Young and R Jaenisch. (2006). Polycomb complexes repress developmental regulators in murine embryonic stem cells. Nature 441:349–53.

10. Pasini D, AP Bracken, JB Hansen, M Capillo and K Helin. (2007). The polycomb group protein Suz12 is required for embryonic stem cell differentiation. Mol Cell Biol 27:3769–79.

11. Schuettengruber B, HM Bourbon, L Di Croce and G Cavalli. (2017). Genome Regulation by Polycomb and Trithorax: 70 Years and Counting. Cell 171:34–57.

12. Muller S and A Nayak. (2016). Inhibition of MLL1 histone methyltransferase brings the developmental clock back to naive pluripotency. Stem Cell Investig 3:58.

13. Zhang H, S Gayen, J Xiong, B Zhou, AK Shanmugam, Y Sun, H Karatas, L Liu, RC Rao, S Wang, AI Nesvizhskii, S Kalantry and Y Dou. (2016). MLL1 Inhibition Reprograms Epiblast Stem Cells to Naive Pluripotency. Cell Stem Cell 18:481–94.

14. Zhang H, LTP Khoa, F Mao, H Xu, B Zhou, Y Han, M O’Leary, A Nusrat, L Wang, TL Saunders and Y Dou. (2019). MLL1 Inhibition and Vitamin D Signaling Cooperate to Facilitate the Expanded Pluripotency State. Cell Rep 29:2659–2671 e6.

15. Tammimies K, CR Marshall, S Walker, G Kaur, B Thiruvahindrapuram, AC Lionel, RK Yuen, M Uddin, W Roberts, R Weksberg, M Woodbury-Smith, L Zwaigenbaum, E Anagnostou, Z Wang, J Wei, JL Howe, MJ Gazzellone, L Lau, WW Sung, K Whitten, C Vardy, V Crosbie, B Tsang, L D’Abate, WW Tong, S Luscombe, T Doyle, MT Carter, P Szatmari, S Stuckless, D Merico, DJ Stavropoulos, SW Scherer and BA Fernandez. (2015). Molecular Diagnostic Yield of Chromosomal Microarray Analysis and Whole-Exome Sequencing in Children With Autism Spectrum Disorder. JAMA 314:895–903.

16. Wang T, H Guo, B Xiong, HA Stessman, H Wu, BP Coe, TN Turner, Y Liu, W Zhao, K Hoekzema, L Vives, L Xia, M Tang, J Ou, B Chen, Y Shen, G Xun, M Long, J Lin, ZN Kronenberg, Y Peng, T Bai, H Li, X Ke, Z Hu, J Zhao, X Zou, K Xia and EE Eichler. (2016). De novo genic mutations among a Chinese autism spectrum disorder cohort. Nat Commun 7:13316.

17. Stessman HA, B Xiong, BP Coe, T Wang, K Hoekzema, M Fenckova, M Kvarnung, J Gerdts, S Trinh, N Cosemans, L Vives, J Lin, TN Turner, G Santen, C Ruivenkamp, M Kriek, A van Haeringen, E Aten, K Friend, J Liebelt, C Barnett, E Haan, M Shaw, J Gecz, BM Anderlid, A Nordgren, A Lindstrand, C Schwartz, RF Kooy, G Vandeweyer, C Helsmoortel, C Romano, A Alberti, M Vinci, E Avola, S Giusto, E Courchesne, T Pramparo, K Pierce, S Nalabolu, DG Amaral, IE Scheffer, MB Delatycki, PJ Lockhart, F Hormozdiari, B Harich, A Castells-Nobau, K Xia, H Peeters, M Nordenskjold, A Schenck, RA Bernier and EE Eichler. (2017). Targeted sequencing identifies 91 neurodevelopmental-disorder risk genes with autism and developmental-disability biases. Nat Genet 49:515–526.

18. Zhu L, Q Li, SH Wong, M Huang, BJ Klein, J Shen, L Ikenouye, M Onishi, D Schneidawind, C Buechele, L Hansen, J Duque-Afonso, F Zhu, GM Martin, O Gozani, R Majeti, TG Kutateladze and ML Cleary. (2016). ASH1L Links Histone H3 Lysine 36 Dimethylation to MLL Leukemia. Cancer Discov 6:770–83.

19. Tanaka Y, K Kawahashi, Z Katagiri, Y Nakayama, M Mahajan and D Kioussis. (2011). Dual function of histone H3 lysine 36 methyltransferase ASH1 in regulation of Hox gene expression. PLoS One 6:e28171.

20. Tanaka Y, Z Katagiri, K Kawahashi, D Kioussis and S Kitajima. (2007). Trithorax-group protein ASH1 methylates histone H3 lysine 36. Gene 397:161–8.

21. Gregory GD, CR Vakoc, T Rozovskaia, X Zheng, S Patel, T Nakamura, E Canaani and GA Blobel. (2007). Mammalian ASH1L is a histone methyltransferase that occupies the transcribed region of active genes. Mol Cell Biol 27:8466–79.

22. Bertero A, P Madrigal, A Galli, NC Hubner, I Moreno, D Burks, S Brown, RA Pedersen, D Gaffney, S Mendjan, S Pauklin and L Vallier. (2015). Activin/nodal signaling and NANOG orchestrate human embryonic stem cell fate decisions by controlling the H3K4me3 chromatin mark. Genes Dev 29:702–17.

23. Huang C and B Zhu. (2018). Roles of H3K36-specific histone methyltransferases in transcription: antagonizing silencing and safeguarding transcription fidelity. Biophys Rep 4:170–177.

24. Dorighi KM and JW Tamkun. (2013). The trithorax group proteins Kismet and ASH1 promote H3K36 dimethylation to counteract Polycomb group repression in Drosophila. Development 140:4182–92.

25. Miyazaki H, K Higashimoto, Y Yada, TA Endo, J Sharif, T Komori, M Matsuda, Y Koseki, M Nakayama, H Soejima, H Handa, H Koseki, S Hirose and K Nishioka. (2013). Ash1l methylates Lys36 of histone H3 independently of transcriptional elongation to counteract polycomb silencing. PLoS Genet 9:e1003897.

26. Aloia L, B Di Stefano and L Di Croce. (2013). Polycomb complexes in stem cells and embryonic development. Development 140:2525–34.

27. Yin B, F Yu, C Wang, B Li, M Liu and L Ye. (2019). Epigenetic Control of Mesenchymal Stem Cell Fate Decision via Histone Methyltransferase Ash1l. Stem Cells 37:115–127.

28. Jones M, J Chase, M Brinkmeier, J Xu, DN Weinberg, J Schira, A Friedman, S Malek, J Grembecka, T Cierpicki, Y Dou, SA Camper and I Maillard. (2015). Ash1l controls quiescence and self-renewal potential in hematopoietic stem cells. J Clin Invest 125:2007–20.

29. Shi Y, P Kirwan, J Smith, HP Robinson and FJ Livesey. (2012). Human cerebral cortex development from pluripotent stem cells to functional excitatory synapses. Nat Neurosci 15:477–86, S1.

30. Shi Y, P Kirwan and FJ Livesey. (2012). Directed differentiation of human pluripotent stem cells to cerebral cortex neurons and neural networks. Nat Protoc 7:1836–46.

31. Lizarraga SB, L Ma, AM Maguire, LI van Dyck, Q Wu, Q Ouyang, BC Kavanaugh, D Nagda, LL Livi, MF Pescosolido, M Schmidt, S Alabi, MH Cowen, P Brito-Vargas, D Hoffman-Kim, ED Gamsiz Uzun, A Schlessinger, RN Jones and EM Morrow. (2021). Human neurons from Christianson syndrome iPSCs reveal mutation-specific responses to rescue strategies. Sci Transl Med 13.

32. Thomson JA, J Itskovitz-Eldor, SS Shapiro, MA Waknitz, JJ Swiergiel, VS Marshall and JM Jones. (1998). Embryonic stem cell lines derived from human blastocysts. Science 282:1145–7.

33. Guschin DY, AJ Waite, GE Katibah, JC Miller, MC Holmes and EJ Rebar. (2010). A rapid and general assay for monitoring endogenous gene modification. Methods Mol Biol 649:247–56.

34. Findlay SD, KM Vincent, JR Berman and LM Postovit. (2016). A Digital PCR-Based Method for Efficient and Highly Specific Screening of Genome Edited Cells. PLoS One 11:e0153901.

35. Miyaoka Y, JR Berman, SB Cooper, SJ Mayerl, AH Chan, B Zhang, GA Karlin-Neumann and BR Conklin. (2016). Systematic quantification of HDR and NHEJ reveals effects of locus, nuclease, and cell type on genome-editing. Sci Rep 6:23549.

36. Miyaoka Y, AH Chan and BR Conklin. (2016). Using Digital Polymerase Chain Reaction to Detect SingleNucleotide Substitutions Induced by Genome Editing. Cold Spring Harb Protoc 2016:pdb prot086801.

37. Miyaoka Y, SJ Mayerl, AH Chan and BR Conklin. (2018). Detection and Quantification of HDR and NHEJ Induced by Genome Editing at Endogenous Gene Loci Using Droplet Digital PCR. Methods Mol Biol 1768:349–362.

38. Ouyang Q, SB Lizarraga, M Schmidt, U Yang, J Gong, D Ellisor, JA Kauer and EM Morrow. (2013). Christianson syndrome protein NHE6 modulates TrkB endosomal signaling required for neuronal circuit development. Neuron 80:97–112.

39. Brinkman EK, T Chen, M Amendola and B van Steensel. (2014). Easy quantitative assessment of genome editing by sequence trace decomposition. Nucleic Acids Res 42:e168.

40. Dehairs J, A Talebi, Y Cherifi and JV Swinnen. (2016). CRISP-ID: decoding CRISPR mediated indels by Sanger sequencing. Sci Rep 6:28973.

41. Watanabe K, M Ueno, D Kamiya, A Nishiyama, M Matsumura, T Wataya, JB Takahashi, S Nishikawa, S Nishikawa, K Muguruma and Y Sasai. (2007). A ROCK inhibitor permits survival of dissociated human embryonic stem cells. Nat Biotechnol 25:681–6.

42. Ho SM, BJ Hartley, E Flaherty, P Rajarajan, R Abdelaal, I Obiorah, N Barretto, H Muhammad, HP Phatnani, S Akbarian and KJ Brennand. (2017). Evaluating Synthetic Activation and Repression of Neuropsychiatric-Related Genes in hiPSC-Derived NPCs, Neurons, and Astrocytes. Stem Cell Reports 9:615–628.

43. Lino CA, JC Harper, JP Carney and JA Timlin. (2018). Delivering CRISPR: a review of the challenges and approaches. Drug Deliv 25:1234–1257.

44. Brinkman EK and B van Steensel. (2019). Rapid Quantitative Evaluation of CRISPR Genome Editing by TIDE and TIDER. Methods Mol Biol 1961:29–44.

45. Mitsui K, Y Tokuzawa, H Itoh, K Segawa, M Murakami, K Takahashi, M Maruyama, M Maeda and S Yamanaka. (2003). The homeoprotein Nanog is required for maintenance of pluripotency in mouse epiblast and ES cells. Cell 113:631–42.

46. Satterstrom FK, RK Walters, T Singh, EM Wigdor, F Lescai, D Demontis, JA Kosmicki, J Grove, C Stevens, J Bybjerg-Grauholm, M Baekvad-Hansen, DS Palmer, JB Maller, P-BC i, M Nordentoft, O Mors, EB Robinson, DM Hougaard, TM Werge, P Bo Mortensen, BM Neale, AD Borglum and MJ Daly. (2019). Autism spectrum disorder and attention deficit hyperactivity disorder have a similar burden of rare protein-truncating variants. Nat Neurosci 22:1961–1965.

47. Zhang C, L Xu, X Zheng, S Liu and F Che. (2020). Role of Ash1l in Tourette syndrome and other neurodevelopmental disorders. Dev Neurobiol.

48. Liu S, M Tian, F He, J Li, H Xie, W Liu, Y Zhang, R Zhang, M Yi, F Che, X Ma, Y Zheng, H Deng, G Wang, L Chen, X Sun, Y Xu, J Wang, Y Zang, M Han, X Wang, H Guan, Y Ge, C Wu, H Wang, H Liang, H Li, N Ran, Z Yang, H Huang, Y Wei, X Zheng, X Sun, X Feng, L Zheng, T Zhu, W Luo, Q Chen, Y Yan, Z Huang, Z Jing, Y Guo, X Zhang, CP Schaaf, J Xing, C Wang, F Yu and JS Guan. (2020). Mutations in ASH1L confer susceptibility to Tourette syndrome. Mol Psychiatry 25:476–490.

49. Zhu T, C Liang, D Li, M Tian, S Liu, G Gao and JS Guan. (2016). Histone methyltransferase Ash1L mediates activity-dependent repression of neurexin-1alpha. Sci Rep 6:26597.

50. Brinkmeier ML, KA Geister, M Jones, M Waqas, I Maillard and SA Camper. (2015). The Histone Methyltransferase Gene Absent, Small, or Homeotic Discs-1 Like Is Required for Normal Hox Gene Expression and Fertility in Mice. Biol Reprod 93:121.

