## Supplementary Material for "Overcoming the effect of ASH1L haploinsufficiency on stem cells amenability to genome editing and differentiation into the neuronal lineage – a technical report"

### SUPPLEMENTARY MATERIALS

Seon Hye Cheon[1,2,§] , Foster D. Ritchie[1,2,§], Nicholas Marinelli[1,2] , Janay M. Vacharasin[1,2] , Collin Cheatle[1,2] , Mikayla M. McCord[1,2], Kaitlin Cox[1,2], and Sofia B. Lizarraga[1,2,ð]

[1] Department of Biological Sciences, University of South Carolina, Columbia, 29208, USA

[2] Center for childhood neurotherapeutics, Department of Biological Sciences, University of South Carolina, Columbia, 29208, USA

[§] These authors contributed equally to this work

[ð] Corresponding author

**SUPPLEMENTARY MATERIAL**

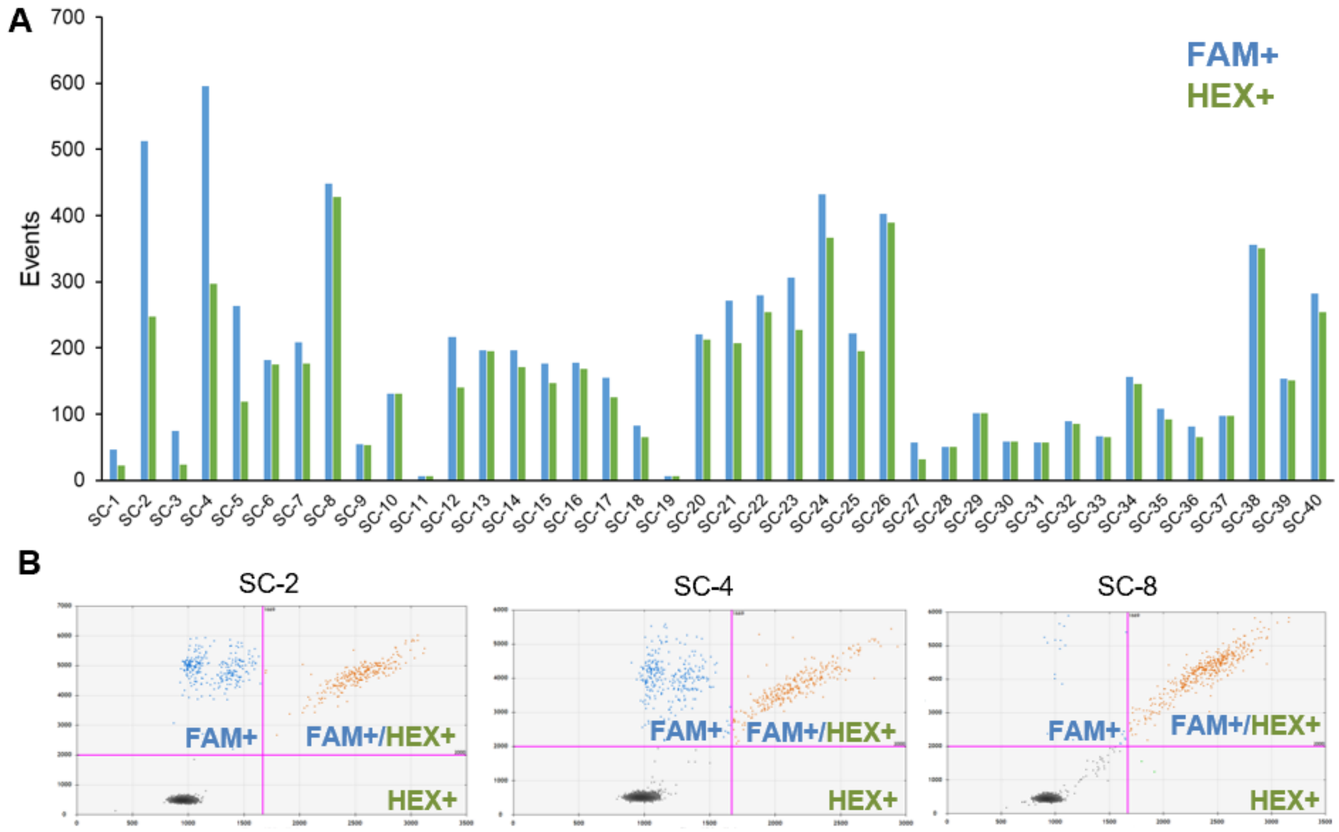

**Supplementary Figure S1. ddPCR Screening of 40 ASH1L Genome-Edited Subclones. (A)** Analysis of 40 subclonal ESC populations after nucleofection with ASH1L-sgRNA set-4 for FAM+ and HEX+ signal. ddPCR was conducted on genomic DNA extracted from each clonal line. **(B)** A 2D plot of droplet fluorescence for FAM and HEX levels is shown for ASH1L mutant clone lines, SC-2 and SC-4, and the un-edited line SC-8.

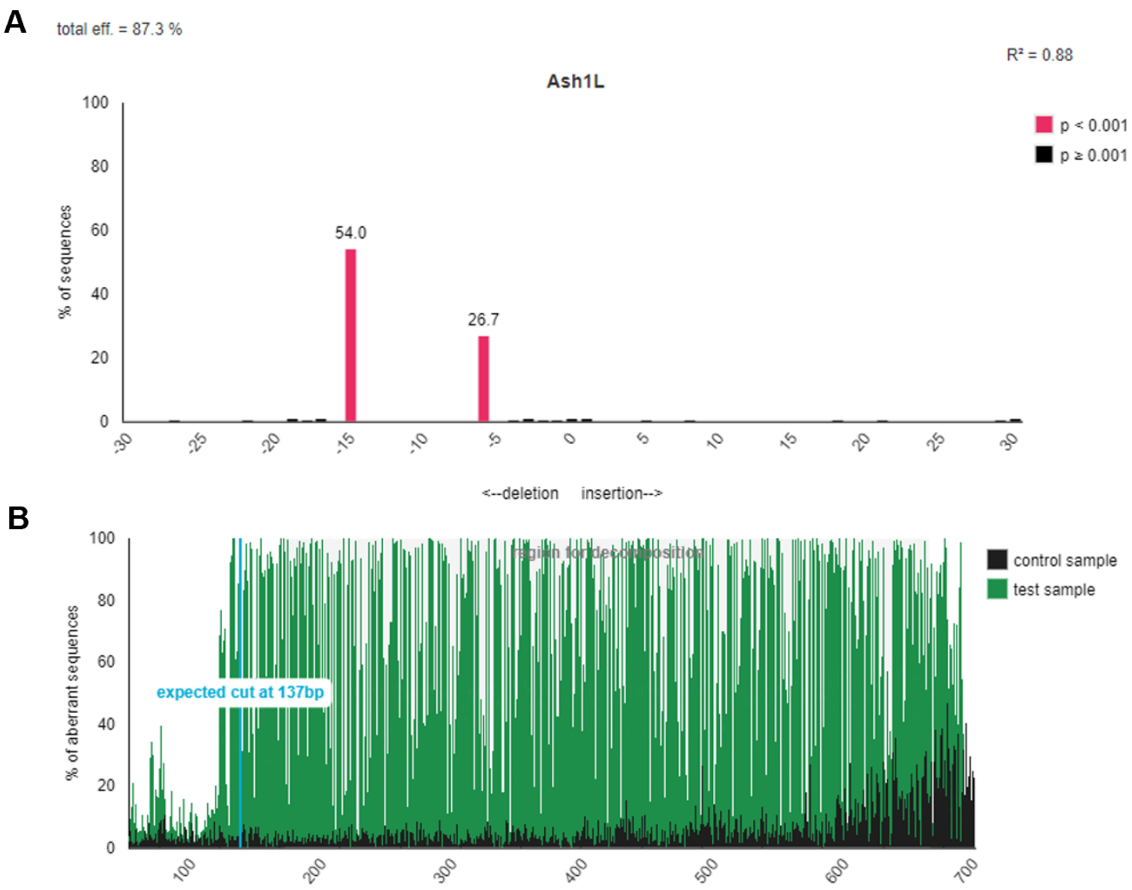

Supplementary

**Supplementary Figure S2. Predicted Indels and Aberrant Signals of ASH1L genome-edited Subclones. (A)** Indels Spectrum of SC-1.1 predicting two deletions at -15bp and -6bp from expected cut site. **(B)** Percentage of aberrant sequences of SC-1.1 indicating an expected cut side at c.1344 bp.

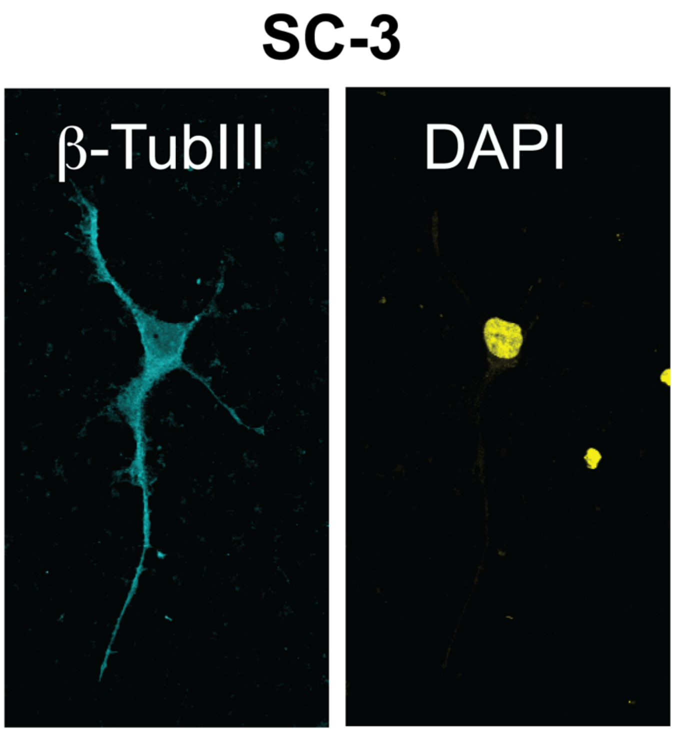

**Supplementary Figure S3. ASH1L haploinsufficiency does not precludes the generation of human neurons.** SC-3 was differentiated using a dual-SMAD inhibition protocol and stained with pan neuronal marker b-tubulin III (cyan) and Dapi (yellow).

| Supplementary Table S1- Cell Culture Media & Buffers | | | |
|---|---|---|---|
| **HEK-cell Media** | **Volume** | **Company** | **Catalog #** |
| DMEM-High Glucose | 440 mL | Thermo Fisher Scientific | 11965118 |
| Fetal Bovine Serum (Heat Inactivated) | 50 mL | VWR | 3100-500G |
| L-glutamine | 5 mL | Thermo Fisher Scientific | 25-030-081 |
| Penicillin/Streptomycin (100X) | 5 mL | Thermo Fisher Scientific | 15-140-122 |
| **N2 Media** | **Volume** | **Company** | **Catalog #** |
| DMEM-F12/GlutaMAX (1X) | 484.7 mL | Gibco | 10565-018 |
| N2 Supplement (5X) | 5 mL | Fisher Scientific | 17-502-048 |
| L-glutamine (200mM) (100X) | 2.5 mL | Thermo Fisher Scientific | 25-030-081 |
| MEM-NEAA (100X) | 5 mL | Thermo Fisher Scientific | 11-140-050 |
| Beta-Mercaptoethanol (14.3M) | 3.49 uL | Life Technologies | 31350-010 |
| Penicillin/Streptomycin (100X) | 2.5 mL | Thermo Fisher Scientific | 15-140-122 |
| Insulin (10mg/mL) | 250 uL | Sigma-Aldrich | I0516-5ML |
| **B-27 Media** | **Volume** | **Company** | **Catalog #** |
| Neurobasal Medium | 482.5 mL | ThermoFisher scientific | 21103049 |
| B-27 Supplement (50X) | 10 mL | ThermoFisher scientific | A3582801 |
| L-glutamine (200mM) (100X) | 5 mL | Thermo Fisher Scientific | 25-030-081 |
| Penicillin/Streptomycin (100X) | 2.5 mL | Thermo Fisher Scientific | 15-140-122 |
| **Neural Induction Medium (NMM)** | **Volume** | **Company** | **Catalog #** |
| B27 media + N2 media (1:1) | 25 mL | N/A | N/A |
| **Neural Induction Medium (NMM)** | **Volume** | **Company** | **Catalog #** |
| Neural Maintenance Medium | 50 mL | N/A | N/A |
| Dorsomorphin (Stock 10mM in DMSO) | 5 $\mu$L | Stemgent | 04-2024 |
| SB431542 (Stock 20mM in DMSO) | 25 $\mu$L | STEMCELL Technologies | 72232 |
| **mTSER Complet Kit - GMP + Pen/Strep** | **Volume** | **Company** | **Catalog #** |
| mTeSR1 basal media + 5X supplement | 500 mL & 50 ml | STEMCELL Technologies | 85850 |
| Penicillin/Streptomycin (100X) | 2.5 mL | Thermo Fisher Scientific | 15-140-122 |
| **KO-DMEM** | **Volume** | **Company** | **Catalog #** |
| KO-DMEM | 49.98 mL | Gibco | 10829018 |
| 20% KO-serum | 10 mL | Gibco | 10828010 |
| 1% Glutamax | 500 $\mu$L | Gibco | 35050079 |
| 1% MEM-NEAA | 500 $\mu$L | Thermo Fisher Scientific | 11-140-050 |
| 0.1% Beta-mercaptoethanol | 50 $\mu$L | Life Technologies | 31350-010 |
| 20ng/mL bFGF | 10 $\mu$L | PeproTech | 100-18C |
| **RIPA Buffer** | **Volume** | **Company** | **Catalog #** |
| 1M NaCl | 15 mL | Sigma | S3014-1KG |
| 1M Tris-HCl pH 8.0 | 5 mL | MP Biomedicals | 103133 |
| 0.5M Na Fluoride | 2 mL | MP Biomedicals | 194864 |
| 100% Nonidet p-40 | 1 mL | Sigma | N-6507 |
| 10% Na Deoxycholate | 5 mL | Sigma-Aldrich | D6750-100G |
| 10% SDS | 1 mL | Thermo Fisher Scientific | 15553027 |
| Protease Inhibitor (1:500) | 200 $\mu$L | Sigma Aldrich | P8340 |
| Phosphatase Inhibitor (1:500) | 200 $\mu$L | Fisher Scientific | PIA32961 |
| **TowBin Buffer, pH 8.3** | **Volume /Weight** | **Company** | **Catalog #** |
| Tris Base (final 20mM) | 3.03 g | MP Biomedicals | 103133 |
| Glycine (final 192mM) | 14.4 g | Sigma | G8898-1KG |
| Methanol (final 20%) | 200 mL | Sigma-Aldrich | 322415-2L |
| **TBST** | **Volume / Weight** | **Company** | **Catalog #** |
| Tris HCl (final 10mM) | 1.57 g | Sigma-Aldrich | 648317-1KG |
| NaCl (final 150mM) | 8.77 g | Sigma-Aldrich | S9625-1KG |
| Tween-20 (final 0.1%) | 1 mL | Sigma-Aldrich | P9416-50ML |

**Supplementary Table S1. Media and Associated Reagents.** Composition of medias and buffers used for cell culture and experiments. Company and catalog number for each reagent is provided.

| Supplementary Table S2- General Reagents For Cellular, Molecular Biology & Biochemistry |
|---|

| hESC Reagents | Company | Catalog # |
|---|---|---|
| Y-27632 | stem cell technologies | 72302 |
| mFreSR | STEMCELL Technologies | 05855 |
| ReLeSR | STEMCELL Technologies | 05872 |
| Gentle Cell Dissociation Reagent (GCDR) | STEMCELL Technologies | 07174 |
| CloneR™ | STEMCELL Technologies | 05888 |
| Puromycin | Life Technologies | A1113803 |
| Corning Matrigel | Fisher Scientific | 08-774-552 |
| Primaria - 6 & 24 multi-well plates | Corning | 353846 / 7 |
| **Transfection Reagents** | **Company** | **Catalog #** |
| P3 Primary Cell 4D-Nucleofector® X Kit S (32 RCT) | Lonza | V4XP-3032 |
| Lipofectamine 2000 Transfection Kit | Thermo Fisher Scientific | 11668030 |
| **Neuronal Differentiation reagents** | **Company** | **Catalog #** |
| Recombinant FGF-basic | PeproTech | 100-18C |
| STEMdiff™ Neural Induction Medium | STEMCELL Technologies | 05835 |
| Laminin | Invitrogen | 23017015 |
| Poly-Ornithine | Sigma-Aldrich | P4957 |
| **Molecular Biology Reagents and Kits** | **Company** | **Catalog #** |
| Qiagen Endofree Maxi Kit | Qiagen | 12362 |
| Qiagen QIAquick Gel Extraction Kit | Qiagen | 28704 |
| DNeasy Blood and Tissue Kit | Qiagen | 69504 |
| Sensifast cDNA Conversion kit | Meridian Bioscience | BIO-65054 |
| RNeasy Micro Kit | Qiagen | 74304 |
| SYBR Green Master Mix | Thermo Fisher Scientific | A46012 |
| Plasmid-Safe™ ATP-Dependent DNase (1mM) | Epicentre | E3101K |
| FastDigest BbsI | Thermo Scientific | FD1014 |
| EcoRI-HF | New England Biolabs | R3101S |
| Stbl3 Competent E. Coli | Invitrogen | C737303 |
| SOC Media | KD Medical Inc | BLA-5140 |
| Ampicillin | Sigma-Aldrich | A9393 |
| LB Agar | Fisher Scientific | BP9745-500 |
| LB Broth Miller Powder | Fisher Scientific | BP1426500 |
| Herculase | Agilent | 600677 |
| Herculase Buffer | Agilent | 600677 |
| Magnesium Chloride | Sigma-Aldrich | M8266 |
| dNTP | Agilent | 200415 |
| UltraPure 10X TAE Buffer | Invitrogen | 15558-026 |
| 4-20% TBE Gel | Fisher Scientific | EC62255BOX |
| T7 Endonuclease 1 | GeneCopoeia | IC005 |
| QuickExtract™ DNA Extraction Solution | Epicentre | QE09050 |
| Ethidium Bromide | Sigma-Aldrich | E1510 |
| Plasmid-Safe™ ATP-Dependent DNase | Epicentre | E3101K |
| SYBR Gold | Fisher Scientific | S11494 |
| **Biochemistry reagents** | **Company** | **Catalog #** |
| Pierce BCA protein assay kit | Thermo Fisher Scientific | PI23225 |
| Bovine Serum Albumin | Sigma | A3059-100G |
| SuperSignal™ West Pico PLUS Chemiluminescent Substrate | Thermo Scientific | 34578 |
| CL-XPosure™ Film, 5 x 7 in. | Thermo Scientific | 34090 |
| **Cell culture consumables** | **Company** | **Catalog #** |
| Primaria plates (6well and 24 well) | Corning | 35846 / 35847 |

**Supplementary Table S2. General reagents for cellular, molecular biology and biochemistry.**

Company and catalog number of reagents used for experiments in this publication.

| Supplementary Table S3 – qPCR primers, antibodies and plasmids | | |
|---|---|---|
| **Gene** | **qPCR primer Sequences** | **GC Content** |
| *ASH1L* | 5'- GAG TCA CTG CCT TCT AAC GAA -3' | 47.6 |
| | 5'- TCT CAG ATG AAG ACC TTT TCC G -3' | 45.5 |
| *NANOG* | 5'- CCT TCT GCG TCA CAC CAT T -3' | 52.6 |
| | 5'- AAC TCT CCA ACA TCC TGA ACC -3' | 47.6 |
| *AFP* | 5'- TCT GCA TGA ATT ATA CAT TGA CCA C -3' | 36 |
| | 5'- AGG AGA TGT GCT GGA TTG TC -3' | 50 |
| *MYOD* | 5'- TGC TGG ACA GGC AGT CTA - 3' | 55.6 |
| | 5'- CTC CGA CGG CAT GAT GG- 3' | 64.7 |
| *GFAP* | 5'- GCT TCA TCT GCT TCC TGT CT- 3' | 50 |
| | 5' - CTG GAG GTT GAG AGG GAC A- 3' | 57.9 |
| *GAPDH* | 5'- ACA TCG CTC AGA CAC CAT G- 3' | 52.6 |
| | 5'- TGT AGT TGA GGT CAA TGA AGG G -3' | 45.5 |
| *EEF2* | 5'- GAC ACG CTT CAC TGA TAC CC -3' | 55 |
| | 5'- TTC AAG TCA TTC TCC GAG AGC -3' | 47.6 |
| **Plasmid** | **Company** | **Catalog #** |
| pSpCas9n(BB)-2A-Puro (Px462) v2 | Addgene | #62987 |
| **Antibodies** | **Company** | **Catalog #** |
| ASH1L | BETHYL Laboratories | A301-748A-T |
| GAPDH | Novus Biologicals | NB600-502 |
| anti-Rabbit Horseradish Peroxidase | Cell Signaling Technology | 7074 |
| Beta Tubulin III | Cell Signaling Technology | 4466 |
| MAP2 | Abcam | AB5392 |

**Supplementary Table S3. qPCR Primers, Plasmids, and Antibodies.** List of primers used for qPCR experiments, along with sequences and GC content for each primer. Identification of plasmid used for transfection. List of antibodies used for western blot and for immunofluorescence analysis. Company and catalog number are also listed for each antibody.

| Supplementary Table S4 – Equipment Used | | |
|---|---|---|
| **Equipment** | **Company** | **Catalog #** |
| 4D-Nucleofector | Lonza | AAF-1002B |
| Thermocycler | Eppendorff | 6321 |
| ChemiDoc MP | Bio-Rad | 12003154 |
| QX200 Droplet Reader | Bio-Rad | 1864003 |
| CFX96 Touch Real Time PCR Detection System | Bio-Rad | 1855195 |
| Spectramax i3 multi-mode platform | Molecular Devices | *Contact manufacturer* |
| CFX384 Optics Module | Bio-Rad | 1855485 |
| SRX-101A Film Processer | Konica | 1051/1052 |

**Supplementary Table S4. Equipment.** List of equipment used for experiments in this publication. Company and catalog number are also provided.

| Supplementary Table S5 - Primers for PCR amplification, sequencing & ddPCR | | | |
|---|---|---|---|
| **Primers for amplifying region targeted by sgRNA Sets 1/2** | | | |
| ***Name*** | ***Direction*** | ***Sequence*** | ***Position in Exon 3*** |
| Primer Set 1 | Forward | 5' CATAGTGATGAGACCATTCCCAGTGAT 3' | 4324 bp |
| | Reverse | 5' AACAGCATCCTTTCCAAATCTATAGCG 3' | 5322 bp |
| Primer Set 2 | Forward | 5' TCTTTTGAGCATGTTTCTCTGATTCCC 3' | 4426 bp |
| | Reverse | 5' CTTATGCTTATATCGCTCTCCAACAGC 3' | 5343 bp |
| **Primers for amplifying region targeted by sgRNA Sets 3/4** | | | |
| ***Name*** | ***Direction*** | ***Sequence*** | ***Position in Exon 3*** |
| Primer Set 3 | Forward | 5' GAAGAAGTCATCCGTCTTCATTCACAG 3' | 1228 bp |
| | Reverse | 5' TGCTTTCTTTATTCTGCCGTACAACAT 3' | 2179 bp |
| Primer Set 4 | Forward | 5' CCCTTCAAAGTTGTACAAGAAAGCAGAT 3' | 1173 bp |
| | Reverse | 5' AAAAGTTTTGCTGGTGGCACTATCTTT 3' | 2145 bp |
| **Primers for Sequencing of ASH1L Exon 3 targeted region** | | | |
| ***Name*** | ***Direction*** | ***Sequence*** | ***Position in Exon 3*** |
| Sequencing Primer | Forward | 5' CAGATGATGTTGCAGCCATTGAA 3' | 1196 bp |
| | Reverse | 5' GTTCCTGGCTTTTTGCCTAAGTT 3' | 1730 bp |
| **Primers for ddPCR FAM/HEX Probes** | | | |
| ***Name*** | ***Sequence*** | | ***Position in Exon 3*** |
| FAM - Reference Probe | 5' CGTCTTCATTCACAGGGAGAA 3' | | 1240 bp |
| HEX - Mutant Probe | 5' TCTTCTTGGTAGCCGGGATCCTGAT 3' | | 1335 bp |

**Supplementary Table S5. Primers for DNA Amplification, Sequencing, and FAM/HEX Probes.**

Table shows primers used for DNA Amplification of regions targeted by sgRNA Sets 1/2 and sgRNA Sets 3/4. Additionally, we show primers used for sequencing of ASH1L Exon 3 targeted regions. Primers designed for mutant and reference probes for ddPCR analysis are shown. The FAM probe was created with the reference primer covering a region outside of the targeted region and the HEX probe was created targeting the region expected to cover potential INDEL
